# Transcriptome Profiling Reveals Inhibitory Effect of Down-regulated ZBTB38 gene on the Transcriptional Regulation of Tumor Cells Proliferation

**DOI:** 10.1101/350223

**Authors:** Jie Chen, Chaofeng Xing, Haosen Wang, Zengmeng Zhang, Daolun Yu, Jie Li, Honglin Li, Jun Li, Yafei Cai

## Abstract

Transcription factor ZBTB38 belongs to the zinc finger protein family and contains the typical BTB domains. Only several predicted BTB domain-containing proteins encoded in the human genome have been functionally characterized. No relevant studies have been reported concerning the effect of down-regulated ZBTB38 gene expression on tumor cells through transcriptome analysis. In the present study, 2,438 differentially expressed genes in ZBTB38^−/−^ SH-SY5Y cells were obtained via high-throughput transcriptome sequencing analysis, 83.5% of which was down-regulated. Furthermore, GO functional clustering and KEGG pathway enrichment analysis of these differentially expressed genes (DEGs) revealed that the knocked-down transcription factor ZBTB38 interacted with p53 and arrested cell cycles to inhibit the proliferation of the tumor cells. Besides, it also significantly down-regulated the expressions of PTEN, a “molecular switch” of the PI3K/Akt signaling pathway, and RB1CC1, the key gene for autophagy initiation, and blocked autophagy to accelerate the apoptosis of tumor cells. ZBTB38^−/−^ SH-SY5Y cells were investigated at the whole transcriptome level and key DEGs were screened in the present study for the first time, providing a theoretical foundation for exploring the molecular mechanism of inhibition of tumor cell proliferation and targeted anti-tumor therapies.

## Introduction

The zinc finger and BTB domain-containing protein family (ZBTB) is a class of regulatory proteins that contain multiple C2H2 or C2HC zinc finger domains at the C-terminus and BTB domains at the N-terminus. Most members of the family, as transcription factors, bind to specific DNA sequences and regulate the transcriptional activity of target genes(Stogios et al. 2005; Sasai et al. 2005). In addition, the family members also involve in various intracellular signal transduction pathways via recognizing and interacting with other proteins, thereby playing important roles in the transcriptional repression, DNA damage, tumorigenesis, and cell proliferation, differentiation, and apoptosis(Lee and Maeda 2012; Matsuda et al. 2008; Nishii et al. 2012).

At least 49 ZBTB proteins are encoded in the human genome, most of which are nuclear proteins (Lee and Maeda 2012). The ZBTB38 (also known as CIBZ) belongs to the human zinc finger protein gene family with typical BTB domains. Among the predicted BTB domain-containing proteins encoded by the human genome, only several of them have been functionally characterized (Matsuda et al. 2008; Cai et al. 2012). The present study was designed to analyze the effect of down-regulated expression of ZBTB38 gene on tumor cells in vitro by transcriptome analysis using the neuroblastoma cell model with down-regulated ZBTB38 gene which was successfully established in our previous study s study (Cai et al. 2012; Cai et al. 2017). Consequently, it was found for the first time that when exogenous genes were inserted into the exogenous gene via liposomes, autophagy was not initiated to regulate the protective mechanism of the stress response, which was blocked on the contrary.

Research of the ZBTB protein family is currently focused on the process of tumor formation and infections. However, it is rarely reported with respect to the effect of ZBTB38 on the level of transcriptome of tumor cells. Transcriptomic studies are developing rapidly over recent years, which, contrary to the studies on an individual gene, enable the investigation on the altered expression of differentially expressed genes (DEGs) on the level of whole protein-coding or non-coding RNAs in cells or tissues of the body. Besides, it can also provide information of the relationship between transcriptional regulation and the protein functions in the whole genome under specific conditions (Zhao et al. 2011; Reimann et al. 2014). The development of RNA sequencing (RNA-seq) technology offers important technical support for the annotation and quantification of transcriptomes. The major strength of this technique lies in its high-throughput and high sensitivity for transcript abundance, providing throughout understanding of the transcriptional information of the genome in a comprehensive manner (Chang et al. 2015; Li et al. 2014).

To understand the effect of down-regulated ZBTB38 gene on the transcriptional regulation of tumor cells proliferation, a high-throughput transcriptome sequencing (RNA-seq) approach was adopted to investigate the transcriptome profiles of neuroblastoma cells in which the expression of ZBTB38 gene was down-regulated. Furthermore, DEGs were subjected to bioinformatics analysis of functional clustering with GO annotation and pathway enrichment with KEGG pathway database. Key DEGs were screened and verified by quantitative RT-PCR to obtain more important regulatory genes involved in tumor formation regulated by the transcription factor ZBTB38 and to understand the related regulatory mechanisms, providing a theoretical basis for further exploration of targeted anti-tumor therapies.

## Materials and Methods

### Cell culture and standard assays

SH-SY5Y cells were purchased from American Type Culture Collection (Rockville, MD, USA) and cultured in Dulbecco’s modified Eagle’s medium supplemented with 10% fetal bovine serum and penicillin-streptomycin. Transient transfections, quantitative real-time polymerase chain reaction (qRT-PCR) were performed as described previously (Cai et al. 2012; Cai et al. 2017). The primers used in qRT-PCR and siRNA suppression assays are listed in supplemental Table S1.

### RNA Preparation and library construction for transcriptome sequencing

Transcriptome high-throughput sequencing was performed in the control group (SH-SY5Y cells transfected with liposome alone, Samples-ID: T04, T05, T06) and the treatment group (SH-SY5Y cells transfected with ZBTB38 siRNA, Samples-ID: T01, T02, T07). Total RNA was isolated from SH-SY5Y cells using TRIzol and the pure-link RNA mini kit (ThermoFisher Scientific, Waltham, MA, USA) according to manufacturer’s instructions. RNA purity was checked using the NanoPhotometer spectrophotometer (IMPLEN, CA, USA). RNA concentration was measured using the Qubit RNA Assay Kit in Qubit 2.0 Fluorometer (Life Technologies, CA, USA). RNA integrity was assessed using the RNA Nano 6000 Assay Kit of the Agilent Bioanalyzer 2100 system (Agilent Technologies, CA, USA).

A total amount of 2 p,g RNA per sample was used as input material for the RNA sample preparations. This study included two groups of three biological replicates. Sequencing libraries were generated using NEBNext UltraTM RNA Library Prep Kit for Illumina (NEB, USA) and index codes were added to attribute sequences to each sample. Fragmentation was carried out using divalent cations under elevated temperature in NEBNext First Strand Synthesis Reaction Buffer (5X). First strand cDNA was synthesized using random hexamer primer and M-MuLV Reverse Transcriptase (RNase H-). Second strand cDNA synthesis was subsequently performed using DNA Polymerase I and RNase H. Remaining overhangs were converted into blunt ends via exonuclease/polymerase activities. After adenylation of 3’ ends of DNA fragments, NEBNext Adaptor with hairpin loop structure were ligated to prepare for hybridization. The library fragments were purified with AMPure XP system (Beckman Coulter, Beverly, USA). Then PCR was performed with Phusion High-Fidelity DNA polymerase, Universal PCR primers and Index (X) Primer. At last, PCR products were purified (AMPure XP system) and library quality was assessed on the Agilent Bioanalyzer 2100 system. The clustering of the index-coded samples was performed on a cBot Cluster Generation System using the TruSeq PE Cluster Kit v4-cBot-HS (Illumia). Following cluster generation, the library preparations were sequenced on an Illumina Hiseq 2500 platform, and paired-end reads were generated.

### 4.3 Data and Statistical Analysis

#### 4.3.1 Quality control

Raw reads of fastq format were firstly processed through in-house perl scripts. In this step, clean reads were obtained by removing reads containing adapter, reads containing ploy-N and low quality reads from raw reads. At the same time, Q20, Q30, GC-content and sequence duplication level of the clean reads were calculated. All the downstream analyses were based on clean reads with high quality (Ewing and Green 1998; Ewing et al. 1998). The clean data of this article are publicly available in the NCBI Sequence Reads Archive (SRA) with accession number SRP150042.

#### 4.3.2 Comparative analysis

The adaptor sequences and low-quality sequence reads were removed from the data sets. Raw sequences were transformed into clean reads after data processing. These clean reads were then mapped to the reference genome sequence. Only reads with a perfect match or one mismatch were further analyzed and annotated based on the reference genome. Tophat2 tools soft were used to map with reference genome (Langmead et al. 2009; Kim et al. 2013). Reference genome download address: ftp://ftp.ensembl.org/pub/release-80/fasta/homosapiens/.

#### 4.3.3 Gene functional annotation

The assembled sequences were compared against the NR(NCBI non-redundant protein sequences), Nt (NCBI non-redundant nucleotide sequences), Pfam (<u>Protein family</u>), Swiss-Prot, KOG/COG (<u>Clusters of Orthologous Groups of proteins</u>), Swiss-Prot (<u>A manually annotated and reviewed protein sequence database</u>), KO (<u>KEGG Ortholog database</u>), and GO (<u>Gene Ontology</u>) databases with an E-value ≤10^−5^ for the functional annotation. The Blast2GO program was used to obtain GO annotation of unigenes including molecular function, biological process, and cellular component categories (Gotz et al. 2008).

#### 4.3.4 Differential expression analysis

Differential expression analysis of the two conditions was performed using the DEGseq R package(Robinson et al. 2010). The P-values obtained from a negative binomial model of gene expression were adjusted using Benjamini and Hochberg corrections to control for false discovery rates (Anders and Huber 2010). Genes with an adjusted P-value < 0.05 were considered to be differently expressed between groups. DEG expression levels were estimated by fragments per kilobase of transcript per million fragments mapped(Florea et al. 2013). The formula is shown as follow:

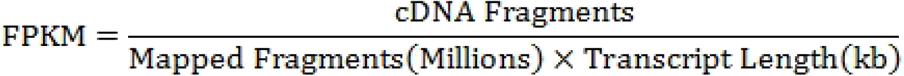

#### 4.3.5 GO enrichment and KEGG pathway enrichment analysis

GO enrichment analysis of the differently expressed genes (DEGs) was implemented in the “GOseq” package in R based on a Wallenius non-central hyper-geometric distribution, which can adjust for gene length bias in DEGs(Young et al. 2010).

KEGG is a database for understanding high-level functions and utilities of biological systems through large-scale molecular datasets generated by genome sequencing and other high-throughput experimental technologies (http://www.genome.jp/kegg/) (Kanehisa et al. 2008). We used the KOBAS software to test for the statistical enrichment of differentially expressed genes in KEGG pathways. KEGG enrichment can identify the principal metabolic pathways and signal transduction pathways of DEGs (Mao et al. 2005).

#### 4.3.6 DEGs quantitative real-time pcr (qRT-PCR) verification

For validation of the transcriptome result, we subjected three significantly differential expressed unigenes on related pathways to qRT-PCR analysis. Redundant RNA from the cDNA library preparation was used to perform reverse transcription according to the Invitrogen protocol. quantitative real-time polymerase chain reaction (qRT-PCR) were performed as described previously (Zhang et al. 2017). The primers used in qRT-PCR suppression assays are listed in Table 1.

#### 4.3.7 Statistical analysis

All data were reported as mean ± standard deviation and analyzed using one-way analysis of variance in SPSS v.17.0. Statistical tests were performed with the Kruskal-Wallis and Mann-Whitney U-tests. A least significant difference test was used for comparisons between groups. A P-value < 0.05 was considered statistically significant.

## 2. Results

### 2.1 Quality control and yield statistics of transcriptome sequencing data

A total of 47.05 Gb clean data were obtained through the transcriptome sequencing of SH-SY5Y cells, with at least 6.12 Gb and a ≥89.30% Q30 percentage for each sample (Table 1). Efficiency of sequence alignment referred to the percentage of mapped reads in the clean reads, which reflected the utilization of transcriptome sequencing data. Statistical analysis of the alignment results showed that the efficiency of read alignment for the reads of each sample and the reference genome ranged between 79.42% and 81.92% (Table 2), which guaranteed that the selected reference genome assembly was qualified for data analysis.

**Table 1.** Summary of Illumina transcriptome sequencing for ZBTB38^−/−^ cells

| Samples-ID | Clean reads | Clean bases | GC Content | %≥Q30 |
| --- | --- | --- | --- | --- |
| T01 | 20,660,153 | 6,116,098,020 | 56.24% | 89.44% |
| T02 | 26,853,046 | 7,955,371,598 | 55.36% | 89.45% |
| T04 | 27,444,964 | 8,143,994,956 | 52.17% | 90.04% |
| T05 | 25,080,118 | 7,422,177,908 | 52.16% | 90.25% |
| T06 | 31,360,838 | 9,309,952,904 | 51.94% | 90.05% |
| T07 | 27,346,767 | 8,098,920,214 | 55.22% | 89.30% |

**Table 2.** Summary of Sequence comparisons among sample sequencing data and selected reference genomes

| Samples-ID | Total Reads | Mapped Reads | Uniq Mapped Reads | Multiple Map Reads |
| --- | --- | --- | --- | --- |
| T01 | 41,320,306 | 32,993,483 (79.85%) | 29,171,906 (70.60%) | 3,821,577 (9.25%) |
| T02 | 53,706,092 | 42,655,511 (79.42%) | 37,610,672 (70.03%) | 5,044,839 (9.39%) |
| T04 | 54,889,928 | 44,964,104 (81.92%) | 41,561,590 (75.72%) | 3,402,514 (6.20%) |
| T05 | 50,160,236 | 40,559,313 (80.86%) | 37,552,835 (74.87%) | 3,006,478 (5.99%) |
| T06 | 62,721,676 | 50,526,963 (80.56%) | 47,347,475 (75.49%) | 3,179,488 (5.07%) |
| T07 | 54,693,534 | 43,544,178 (79.61%) | 38,543,679 (70.47%) | 5,000,499 (9.14%) |

Following the counting of mapped reads at different regions (exons, introns, and intergenic regions) of a given reference genome, distribution maps of mapped reads in different regions of the genome were plotted for each sample. Most reads were mapped to the exon regions of the reference transcriptome (≥80% for each) and the alignment results were valid and reliable (see Figure. S1).

Qualified transcriptome libraries are a major requisite for transcriptome sequencing. To ensure the quality of the libraries, quality of the transcriptome sequencing libraries was evaluated from three different perspectives:

- Randomicity of mRNA fragmentation and the degradation of mRNA were evaluated by examining the distribution of inserted fragments in genes. As shown in Figure S2, the degradation of mRNAs was determined by observing the distribution of mapped reads on mRNA transcripts. The degradation of mRNAs was relatively low in the 6 groups of samples.
- Length of the inserted fragments in the transcripts. The length of the inserted fragment was calculated by the distance between the starting and ending sites of the reads flanking the inserted fragment in the reference genome. Due to the fact that no intron regions were available in the transcriptome-sequenced mRNAs, mRNAs in the transcriptome library were more mature if a single peak on the right side of the main peak was noted in the simulated length distribution map of the inserted fragment. By contrast, the alignment length was longer if more interfering peaks appeared showing the intron regions in the inserted fragment. The dispersion degree of the inserted fragment length directly reflected the efficiency of magnetic bead purification during library preparation. Simulated distribution of the inserted fragment length for each sample showed only single-peak pattern, indicating a high purification rate (see Figure. S3).
- Sequencing saturation status of DEGs in the mapped data was simulated and plotted for the 6 groups of samples, as graphed in the following map. With the increase of sequencing data, the number of DEGs tended to saturate, as shown in Figure S4, which confirmed that the data were sufficient and qualified for the subsequent analysis.

### 2.2 DEG and DEGs Function annotation

To acquire the comprehensive genetic information of ZBTB38^−/−^ SH-SY5Y cells, the unigenes were blasted against the NR, Swiss-Prot, GO, COG, KOG, Pfam, KEGG database resources to identity the functions of all of the unigene sequences. All of DEGs were annotated to genes having known functions in the indicated databases based on the sequences with the greatest similarity. DEseq was used to analyze the DEGs derived from the two groups of cells to obtain a DEGs set. Finally, a total of 2,036 (83.5%) down-regulated DEGs and 402 (16.5%) upregulated DEGs were selected. The number of DEGs annotated in this gene set was shown in Table 3.

**Table 3.**
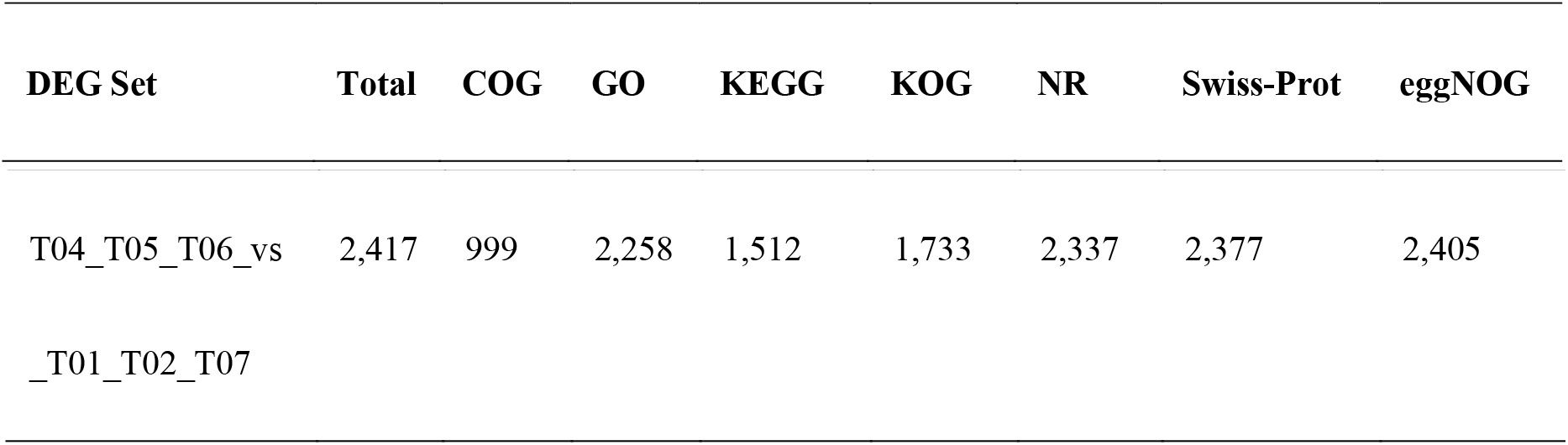
Summary of the function annotation results for ZBTB38^−/−^ unigenes in public protein databases.

A total of 2,258 (93.4%) DEGs were annotated successfully by GO annotation. These annotated DEGs were classified into the next terms of three ontologies: BP (biological process), CC (cellular component) and MF (molecular function). The distribution of unigenes is shown in Figure 1. Among the “Biological Process”, a high percentage of genes were classified into “Cellular Process” (1,924 unigenes, 85.2%), “Single-organism Process” (1827 unigenes, 80.9%) and “biological regulation” (1,786 unigenes, 79.1%). Within the cellular component category, the majority of genes were assigned into “Cell Part” (2,145 unigenes, 95%), “Cell” (2,135 unigenes, 94.6%) and “organelle” (2,023 unigenes, 89.6%). For the molecular function, most of genes were involved in “Binding” (1,949 unigenes, 86.3%) and “Catalytic Activity” (997 unigenes, 44.2%). The greatest number of annotated unigenes were involved in “Biological Process”.

**Figure 1.**
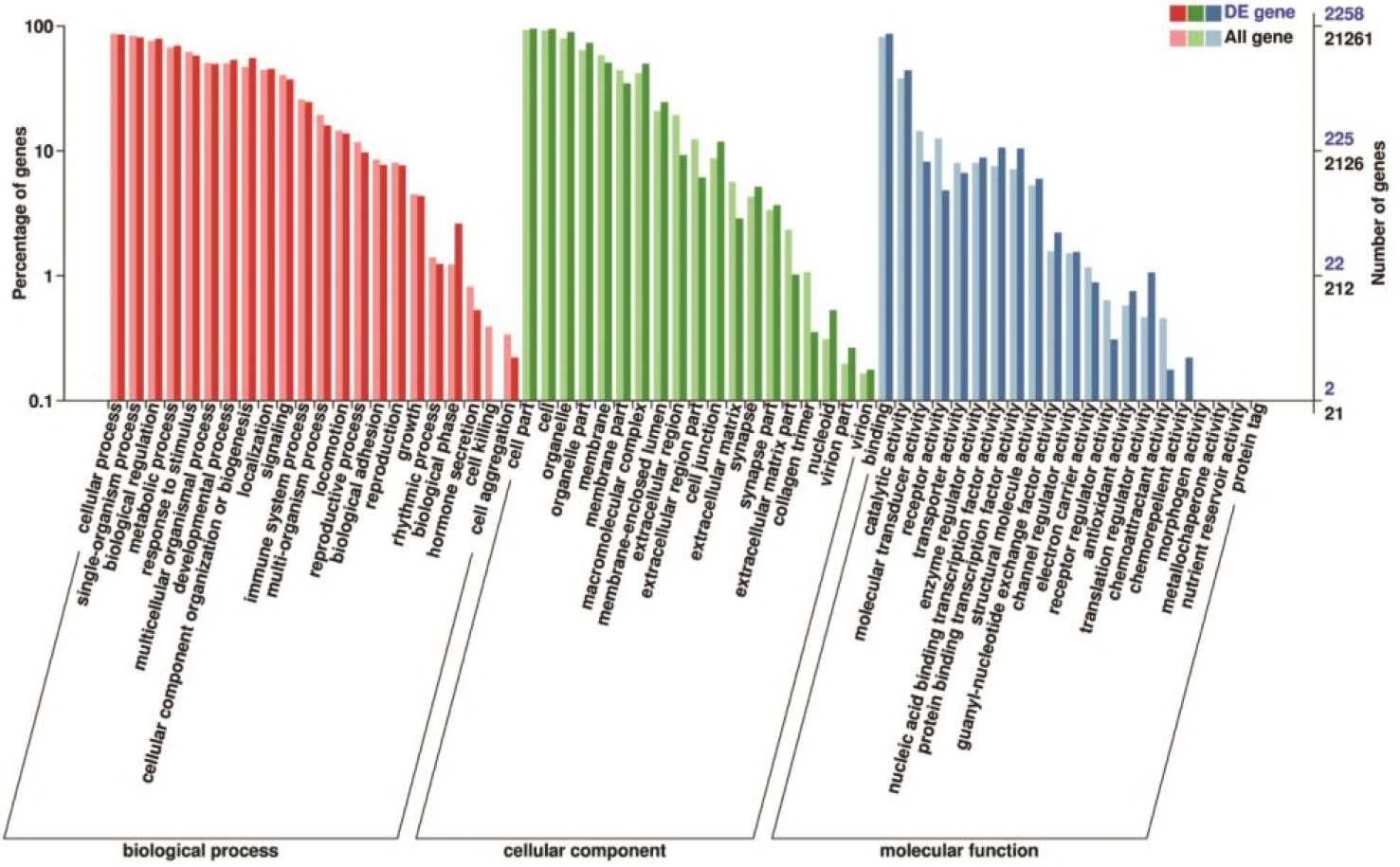
Gene function classification of all annotated unigenes by Gene Ontology. The vertical axis represents the number of unigenes, and horizontal axis gives the specific GO sub-categories.

The unigenes was blasted against the COG database in order to orthologously classify gene products. COG classification statistical results of DEGs were shown in Figure 1. In addition to “General function prediction only”, “Replication, recombination and repair” accounted for the largest proportion of unigenes(180 DEGs, 13.06%), followed by “Transcription” (133 DEGs, 9.65%), “Signal transduction mechanisms”(128 DEGs, 9.29%), “Translation, ribosomal structure and biogenesis” (80 DEGs, 5.81%), “Posttranslational Modification, Protein Turnover and Chaperones” (79 DEGs, 5.73%), “cell cycle control, cell division, and chromosome partitioning” (44 DEGs, 2.98%).

**Figure 2.**
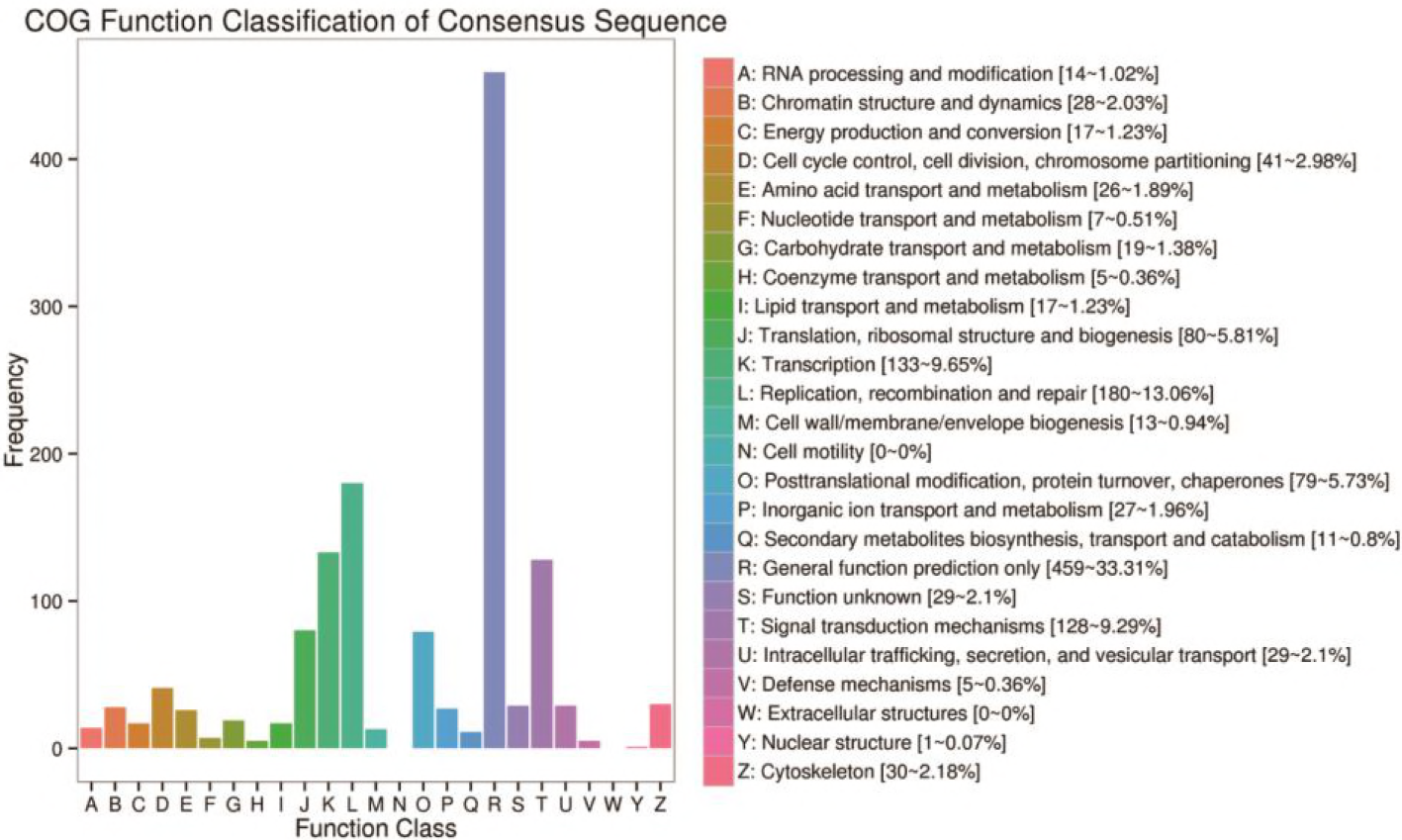
COG function classification of consensus sequence. The vertical axis represents the frequency of unigenes classified into the specific categories, and horizontal axis gives the COG function classification

According to the annotation results of the DEGs KEGG database, KEGG pathways were divided into six branches: “Cellular Processes”, “Environmental Information Processing”, “Genetic Information Processing”, “Human Disease”, “Metabolism” and “Organismal Systems”(Figure 3). Among the KEGG categories, the largest proportion of the unigenes were involved in the “MAPK signaling pathway” and “PI3K-Akt signaling pathway” of “Environmental Information Processing”, “Regulation of actin cytoskeleton” of “Cellular Processes”, and “Ribosomeand” of “Genetic Information Processing”.

**Figure 3.**
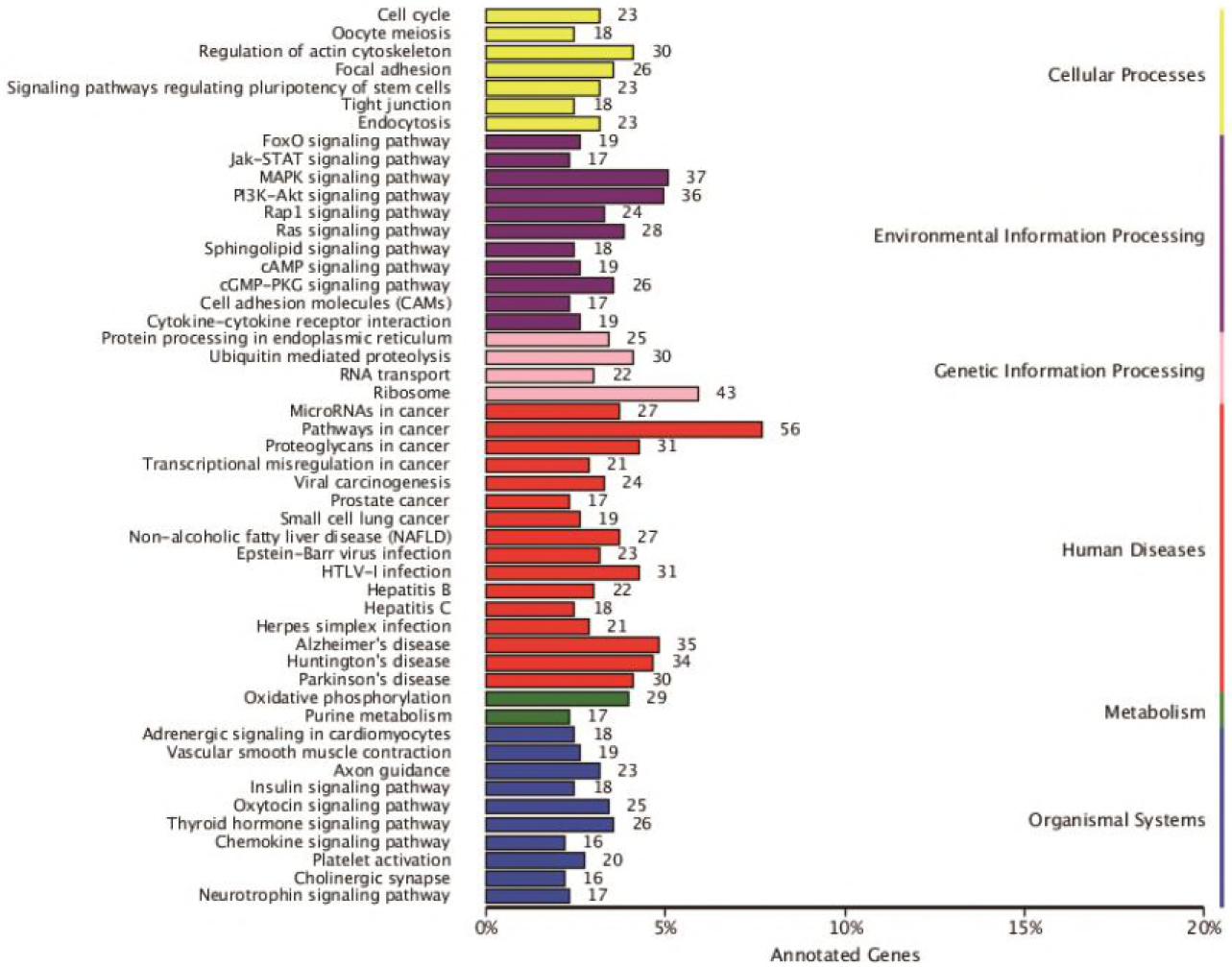
DEG KEGG classification. The vertical axis lists the various metabolic pathways, and horizontal axis gives the percent of annotated genes in the pathways.

Based on the results above, a large number of DEGs were screened after a comparative analysis of relevant databases. Meanwhile, functional annotation was also carried out that was crucial for the further understanding of the cellular functions of ZBTB38 gene as a transcription factor.

### 2.3 DEG KEGG Pathway enrichment and Detection of candidate genes

Pathway enrichment analysis of these DEGs clearly revealed the occurrence of over-presentation of an individual DEG in a given pathway. Pathway enrichment analysis for over-presentation DEGs was performed using the hypergeometric test based on the pathways in the KEGG database to identify pathways that were significantly enriched for DEGs compared to the whole-genome background signals. The results of the KEGG pathway enrichment analysis of DEGs were presented in the figure below, showing the top 20 pathways with the smallest significant q-values (Figure 4). Generally, the largest number of DEGs was enriched in the pathway of “Ribosome”, 39 DEGs were up-regulated and 1 DEG was down-regulated. The results indirectly indicated that in the ZBTB38^−/−^ cells, the function of protein synthesis increased significantly.

**Figure 4.**
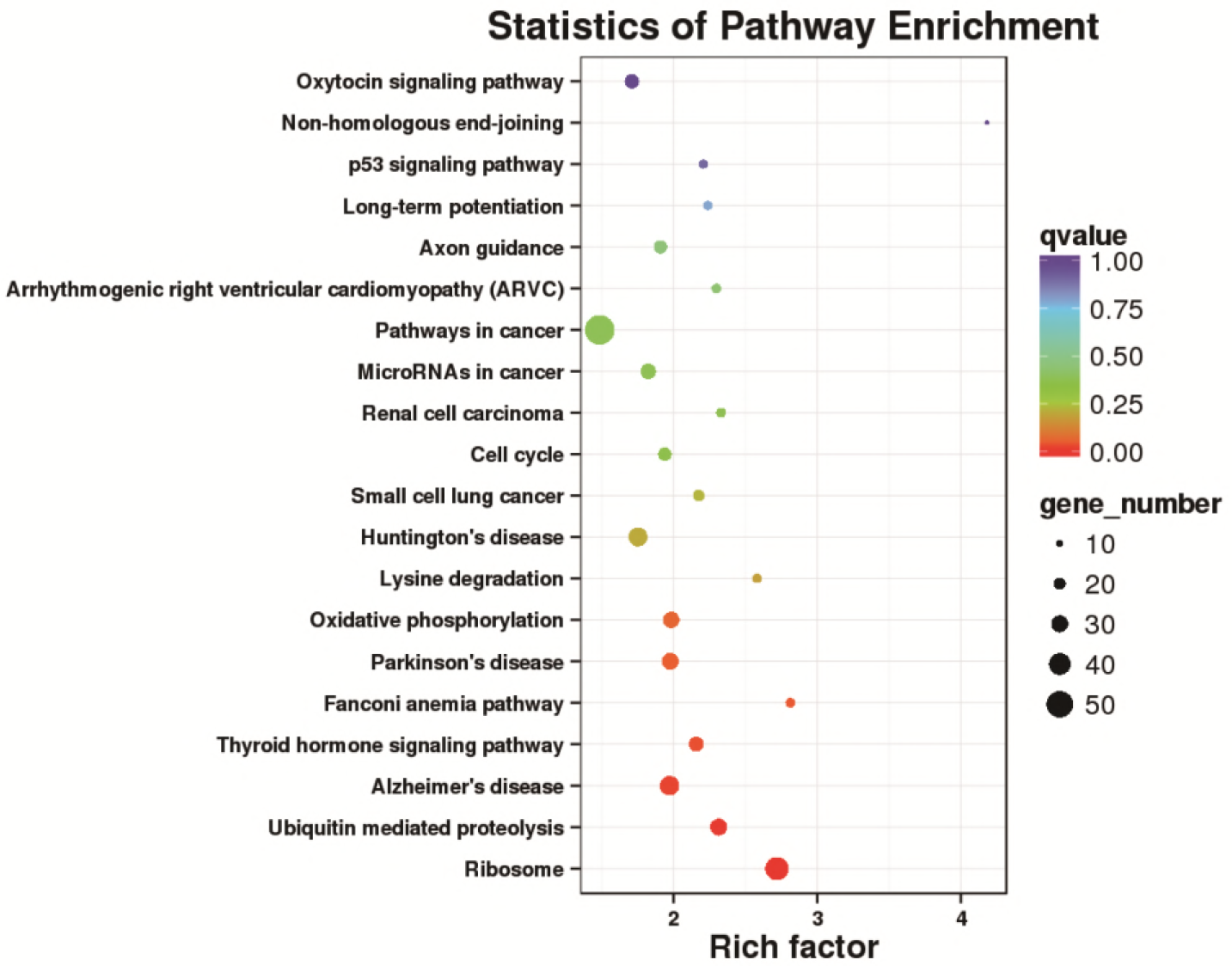
Scatter plot of the KEGG pathway enrichment analysis for DEGs. Each circle in the graph represented a KEGG pathway, with the name of the pathway in the Y-axis and the enrichment factor in the X-axis. An enrichment factor was calculated as the ratio of the proportion of DEGs annotated to a given pathway in all DEGs to the proportion of all genes annotated to the same pathway in all genes. A greater enrichment factor indicted a more significant enrichment of the DEGs in a given pathway. The color of the circle represented the q-value which was the adjusted P after multiple hypothesis testing. A smaller q-value indicated a higher reliability of the significance of the enrichment of DEGs in a given pathway. The size of the circle indicated the number of genes enriched in the pathway, with bigger circle for more genes enriched.

In the KEGG Pathway enrichment analysis, quantification of transcripts and gene expression levels were achieved by Cuffquant and Cuffnorm programs from the Cufflinks software package using the location of a given mapped read on these genes for the 6 groups of samples. Taking FPKM as a measure for the level of transcripts or gene expressions, top 20 down-regulated unigenes associated with autophagy were selected ( see Table S2), among which PIK3C2A was the most down-regulated one, followed by RB1CC1 gene. Genes, including RB1CC1, DDR2, ATM, and FRK, were related to the mTOR signaling pathway and located in the downstream of PI3K-Akt signaling pathway. In summary, the transcription factor ZBTB38 is involved in the process of protein synthesis and also, as a positive regulatory factor, in the occurrence of autophagy directly.

### 2.4 Analysis of the results of Real-time quantitative PCR

To validate the sequencing results obtained by RNA-seq, real-time quantitative PCR was performed on three candidate genes, including PIK3C2A, RB1CC1, ATM, related to the mTOR signaling pathway. The result showed that the expression of these candidate genes was significantly decreased in the ZBTB38^−/−^ cells compared to control group, which was similar to the RNA-seq data (Figure 5). The result verified the reliability of the transcription sequencing results.

**Figure 5.**
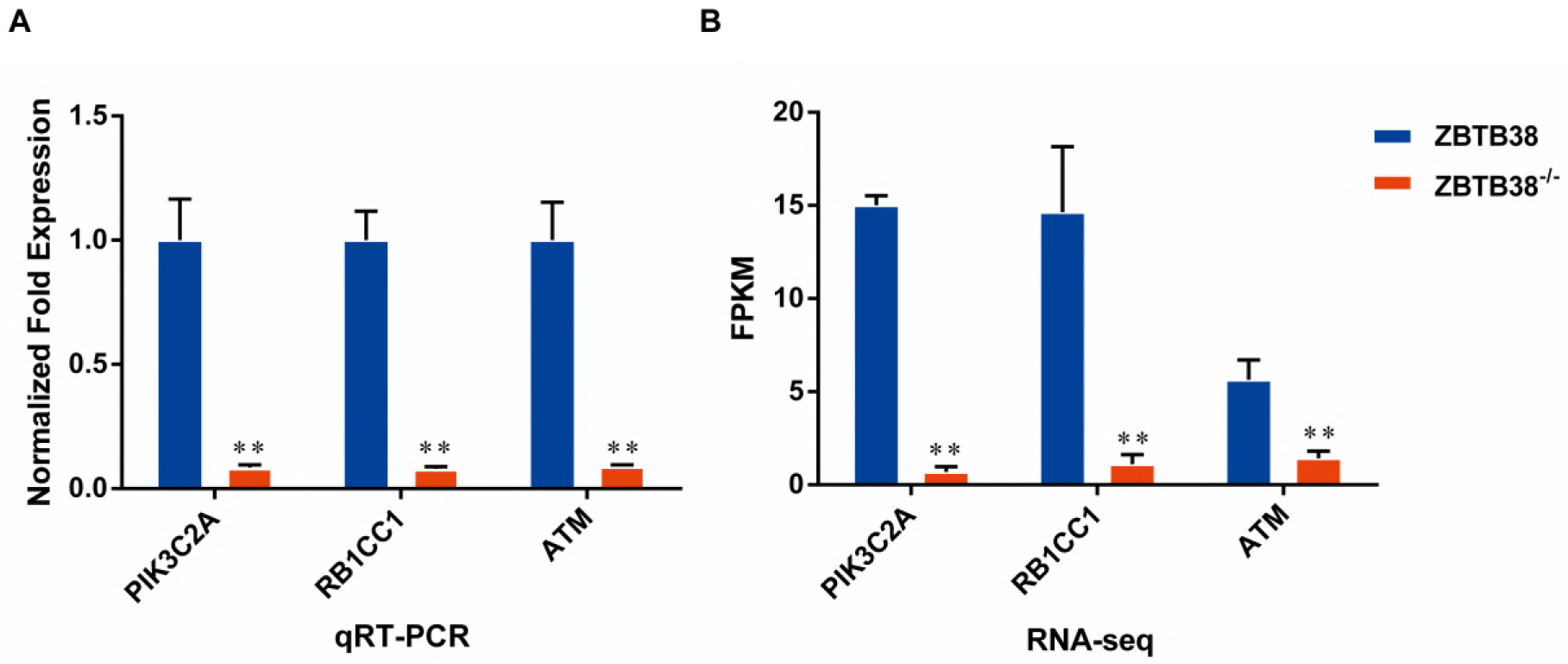
Differential expression analysis of candidate genes between ZBTB38 knockdown (ZBTB38^−/−^) and wild-type SH-SY5Y cells (ZBTB38). (A) The result of qRT-PCR. (B)The result of RNA-seq.

## 3. Discussion

Transcriptomic studies are developing rapidly over recent years. Broadly speaking, the transcriptome is defined as the sum of genes expressed by a single cell of cells group under certain conditions, including the mRNAs, rRNAs, miRNAs, and ncRNAs. In a narrow sense, the transcriptome refers only to the protein-coding mRNAs. The level and pattern of gene expression are different in the body at different growth environment and growth stage, with specific temporal and spatial features, and regulated by both endogenous and exogenous factors. Based on the information of the whole mRNAs obtained in one cell or tissue, transcriptomic studies provide data on the expression regulation systems and protein functions of all genes. Simultaneously, the development of RNA-seq technique offers important technical support for the annotation and quantification of the transcriptome. This research method based on the whole genome of organisms has completely transformed the approach that single-gene study works (Wang et al. 2009; Young et al. 2010; McGettigan 2013).

Among the predicted BTB domain-containing proteins encoded in the human genome, only several of them have been functionally characterized (Lee and Maeda 2012). In the present study, the KEGG pathway enrichment analysis of DEGs revealed that genes in the p53 signaling pathway, including CDK4/6, Cyclin E, MDM2, ATM, ATR, PTEN, were down-regulated, and Gadd45 and PIGs were up-regulated after the knockdown of ZBTB38 (Figure 6). Both the CDK4/6-Cyclin D and the CDK2-Cyclin E complexes are the central links in cell cycle regulation via regulating the G1-S transitions in cells, and abnormal activation of the CyclinD-CDK4/6-INK4-Rb pathway, which is often observed in various malignancies, will lead to uncontrolled growth of cancer cells (The et al. 2015; Sawai et al. 2012; VanArsdale et al. 2015). In addition, members of the Gadd45 family serve as key regulatory genes in DNA damage repair pathway with p53 as the central link, and the upregulation of Gadd45 plays an important role in the regulation of G2/M cell cycle checkpoints and the maintenance of genomic stability, therefore to inhibit the cell transformation and the malignant progression of tumors (Wang et al. 2009). ATM and ATR belong to the inositol trisphosphate kinase family, both of which can be activated by DNA damage to phosphorylate the downstream substrates such as CHK1, CHK2, and p53. In addition, the down-regulation of both kinases may impair the downstream transmission of the molecular signals and inhibit the p53 activity (Matsuoka et al. 2007; Abraham 2001). MDM2 regulates the function of p53 via two approaches, i.e., mediating p53 degradation and inhibiting its transcriptional activity. As a negative feedback regulator of p53, the inhibited expression of MDM2 can enhance the transcriptional activity of p53 and inhibit tumorigenesis (Shangary and Wang 2009). PIGs is a target downstream gene of p53 for the regulation of apoptosis, which is critical for cell apoptosis by participating in the synthesis of reactive oxygen species and the regulation of oxidative stress (Jin et al. 2017; Lee et al. 2010). When ZBTB38 gene is knocked down, p53 expression is decreased and more PIGs are transferred into the nucleus where cell damage is repaired. Therefore, cellular response to DNA damage is increased and the p53-induced ROS production is reduced, eventually leading to the survival of cells. PTEN is a tumor suppressor gene with phosphatase activity. It is an upstream regulatory inhibitor of the PI3K/Akt signal transduction pathway. PTEN is often referred to as a “switch” molecule in the PI3K/Akt pathway due to its ability, which depends on its lipid phosphatase activity, to remove the phosphate group and participate in the regulation of cell activity. Once the expression of PTEN protein is reduced, the dephosphorylation of PIP3 is decreased. Excessive PIP3 is subsequently accumulated in the cells and the PI3K/Akt signaling pathway is continuously activated, eventually leading to cell proliferation or uncontrolled apoptosis and finally the occurrence of various diseases (Bleau et al. 2009; Carnero et al. 2008). Such interpretation may also explain the findings of the present study that DEGs are mainly enriched in the “PI3K-Akt signaling pathway” of the “Environmental Information Processing” category. In summary, direct effect of down-regulated expression of ZBTB38 on the p53 signaling pathway may provide a novel treatment strategy for targeted anti-tumor therapies.

**Figure 6.**
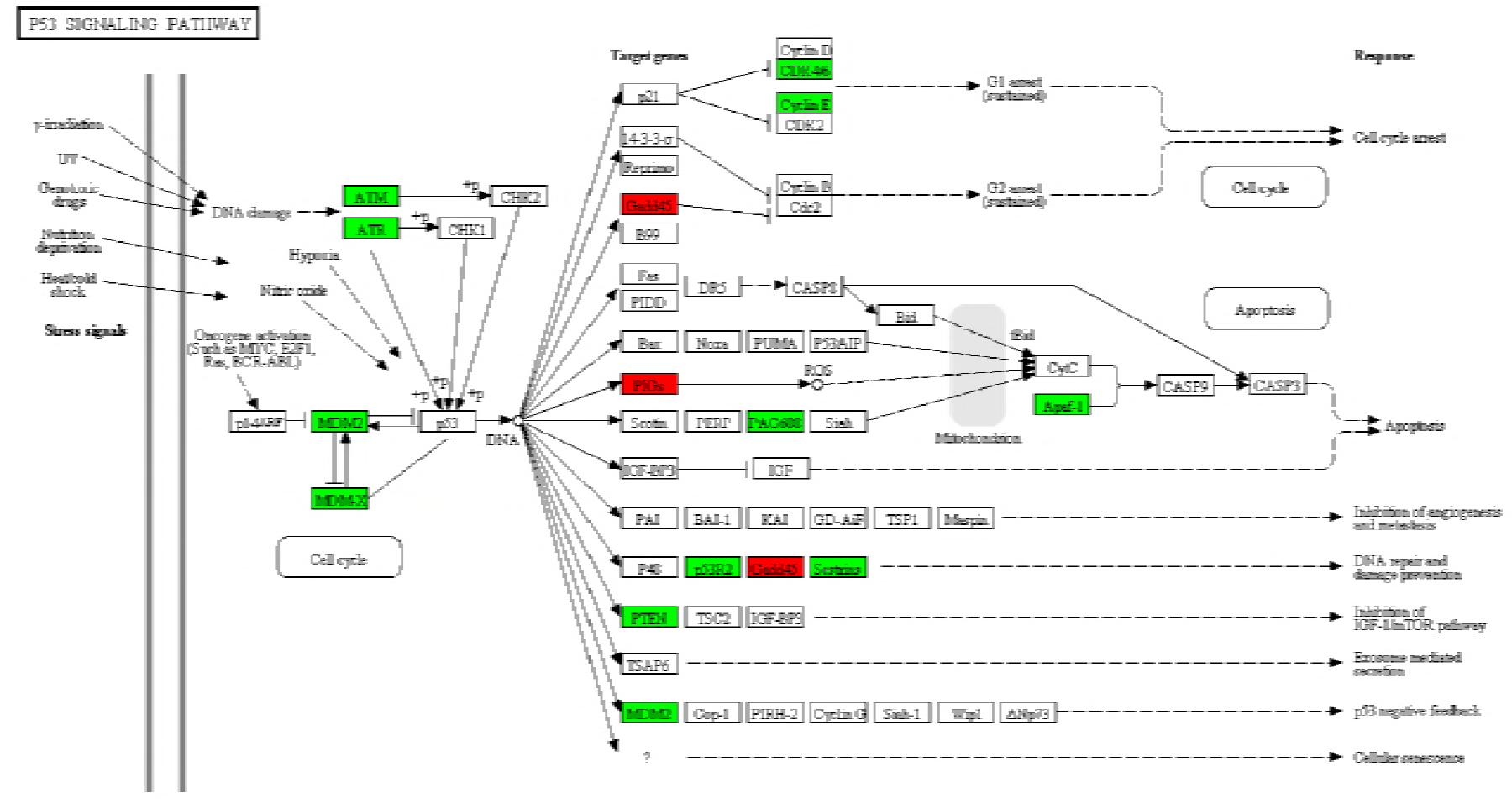
KEGG pathway annotation map of differentially expressed genes in p53 signaling pathway. Relative to the control group, the red box labeled protein was associated with the up-regulated gene and the green box labeled protein was associated with the down-regulated gene.

Majority of the DEGs were enriched in the PI3K-Akt signaling pathway among all KEGG pathway enrichment categories, especially for those down-regulated genes, with most significance noted in PIK3C2A and RB1CC1. PIK3C2A is a member of the PI3Ks family and one of the key molecules in the signal transduction pathway of growth factors. It has been reported that the overexpressed PIK3C2A in cells induces the accumulation and assembly of clathrin, which mediates the transport of proteins between cell membranes and the network structure of the Golgi body via regulating the movement of microtubules (Dragoi and Agaisse 2015; Shi et al. 2016). RB1CC1 (also known as FIP200) is an interacting protein of the focal adhesion kinase family, with a molecular weight of 200kD. As documented in prior studies, autophagy induction is abolished in RB1CC1-deficient cells. RB1CC1 is an important regulatory protein that can acts on the autophagic initiation complex along with the ULK1 simultaneously. Besides, it is also a key autophagy initiation factor in the mTORC1-dependent signaling pathway (Wei et al. 2009; Wang et al. 2011; Ganley et al. 2009). In addition, significant down-regulation of ATM and FOXO1 genes in the ZBTB38^−/−^ cells was observed by transcriptome sequencing analysis in this study. Ataxia telangiectasia mutated (ATM) belongs to the PIKK (PI3K-related protein kinase) family with C-terminal sequences homologous to that of the catalytic region of PI3K. ATM stimulates the downstream signals of the LBK/AMPK/TSC2 pathway and inhibits the mTORC1 to promote cell autophagy (Alexander et al. 2010; Liu et al. 2013). Members of the forkhead box O (FOXO) family are a group of highly conserved transcription factors that play an important role in the regulation of autophagy (Leger et al. 2006). Therefore, it is believed in our study that the autophagy regulation mechanism of the mTORC1-dependent signaling pathway is also inhibited after ZBTB38 knockdown.

Orthologous assignments of gene products were carried out using the COG database. Corresponding statistical analysis of the results also indicated that the silencing of ZBTB38 gene affected the homeostasis of the whole cell, and as a transcriptional factor, ZBTB38 regulated the transcription of intracellular proteins and influenced the expression and transport of proteins in the downstream signaling pathways. The GO functional enrichment analyses of DEGs suggested that most of the genes were involved in “Binding” and “Catalytic Activity” of the molecular function between ZBTB38^−/−^ cells and the controls. Therefore, this also partially explained the biological functions of the key candidate genes enriched in KEGG pathway, i.e., all of them were specific binding DNAs or proteins that regulated the transcriptional activity of target genes and involved in various intracellular signaling pathways.

In conclusion, knockdown of the transcription factor ZBTB38 directly interacted with p53 and arrested cell cycles to inhibit the proliferation of tumor cells. In addition, it also significantly down-regulated the expressions of PTEN, the “molecular switch” of the PI3K/Akt signaling pathway, and RB1CC1, the key gene for autophagy initiation, both of which blocked the autophagy and accelerated the apoptosis of tumor cells. By using neuroblastoma cells, a preliminary study was performed on exploiting DEGs, as well as the key GO terms and KEGG pathways enriched by these DEGs in ZBTB38^−/−^ cells, providing a theoretical foundation for further studies on the regulatory mechanism of ZBTB38 and targeted anti-tumor therapies.

## Author Contributions

Yafei Cai and Jun Li conceived and guided this research. Jie Chen and Chaoxing carried out the experiments, performed data analysis and wrote the manuscript. Haosen Wang and Zengmeng Zhang prepared cell samples and performed the experiments. Daolun Yu and Jie Li performed bioinformatics analysis. Honglin Li analyzed data and discussed results with suggestions from Jie Chen. All authors read and approved the final manuscript.

## Acknowledgments

This study was supported by the National Natural Science Foundation of China (NSFC31372207 and 81570094), the Innovation Team of Scientific Research Platform in Anhui Province, a start-up grant from Nanjing Agricultural University (804090).

## References

Abraham, R.T., 2001 Cell cycle checkpoint signaling through the ATM and ATR kinases. Genes Dev 15 (17):2177–2196. doi:10.1101/gad.914401. http://www.ncbi.nlm.nih.gov/pubmed/11544175

Alexander, A., S.L. Cai, J. Kim, A. Nanez, M. Sahin et al., 2010 ATM signals to TSC2 in the cytoplasm to regulate mTORC1 in response to ROS. Proc Natl Acad Sci U S A 107 (9):4153–4158. doi:10.1073/pnas.0913860107. http://www.ncbi.nlm.nih.gov/pubmed/20160076

Anders, S., and W. Huber, 2010 Differential expression analysis for sequence count data. Genome Biol 11 (10):R106. doi:10.1186/gb-2010-11-10-r106. http://www.ncbi.nlm.nih.gov/pubmed/20979621

Bleau, A.M., D. Hambardzumyan, T. Ozawa, E.I. Fomchenko, J.T. Huse et al., 2009 PTEN/PI3K/Akt pathway regulates the side population phenotype and ABCG2 activity in glioma tumor stem-like cells. Cell Stem Cell 4 (3):226–235. doi:10.1016/j.stem.2009.01.007. http://www.ncbi.nlm.nih.gov/pubmed/19265662

Cai, Y., J. Li, S. Yang, L. Ping, Z. Xuan et al., 2012 CIBZ, a Novel BTB Domain-Containing Protein, Is Involved in Mouse Spinal Cord Injury via Mitochondrial Pathway Independent of p53 Gene. PLoS One 7 (3):e33156.doi:10.1371/journal.pone.0033156. https://www.ncbi.nlm.nih.gov/pubmed/22427977

Cai, Y., J. Li, Z. Zhang, J. Chen, Y. Zhu et al., 2017 Zbtb38 is a novel target for spinal cord injury. Oncotarget 8 (28):45356–45366. doi:10.18632/oncotarget.17487. http://www.ncbi.nlm.nih.gov/pubmed/28514761

Carnero, A., C. Blanco-Aparicio, O. Renner, W. Link, and J.F. Leal, 2008 The PTEN/PI3K/AKT signalling pathway in cancer, therapeutic implications. Curr Cancer Drug Targets 8 (3):187–198. http://www.ncbi.nlm.nih.gov/pubmed/18473732http://www.ncbi.nlm.nih.gov/pubmed/18473732

Chang, Z., G. Li, J. Liu, Y. Zhang, C. Ashby et al., 2015 Bridger: a new framework for de novo transcriptome assembly using RNA-seq data. Genome Biol 16:30. doi:10.1186/s13059-015-0596-2. http://www.ncbi.nlm.nih.gov/pubmed/25723335

Dragoi, A.M., and H. Agaisse, 2015 The class II phosphatidylinositol 3-phosphate kinase PIK3C2A promotes Shigella flexneri dissemination through formation of vacuole-like protrusions. Infect Immun 83 (4):1695–1704. doi:10.1128/IAI.03138-14. http://www.ncbi.nlm.nih.gov/pubmed/25667265

Ewing, B., and P. Green, 1998 Base-calling of automated sequencer traces using phred. II. Error probabilities. Genome Res 8 (3):186–194. http://www.ncbi.nlm.nih.gov/pubmed/9521922

Ewing, B., L. Hillier, M.C. Wendl, and P. Green, 1998 Base-calling of automated sequencer traces using phred. I. Accuracy assessment. Genome Res 8 (3):175–185. http://www.ncbi.nlm.nih.gov/pubmed/9521921

Florea, L., L. Song, and S.L. Salzberg, 2013Thousands of exon skipping events differentiate among splicing patterns in sixteen human tissues. F1000Res 2:188. doi:10.12688/f1000research.2-188.v2. http://www.ncbi.nlm.nih.gov/pubmed/24555089

Ganley, I.G., H. Lam du, J. Wang, X. Ding, S. Chen et al., 2009 ULK1.ATG13.FIP200 complex mediates mTOR signaling and is essential for autophagy. J Biol Chem 284 (18):12297–12305. doi:10.1074/jbc.M900573200. http://www.ncbi.nlm.nih.gov/pubmed/19258318

Gotz, S., J.M. Garcia-Gomez, J. Terol, T.D. Williams, S.H. Nagaraj et al., 2008 High-throughput functional annotation and data mining with the Blast2GO suite. Nucleic Acids Res 36 (10):3420–3435. doi:10.1093/nar/gkn176. http://www.ncbi.nlm.nih.gov/pubmed/18445632

Jin, M., S.J. Park, S.W. Kim, H.R. Kim, J.W. Hyun et al., 2017 PIG3 Regulates p53 Stability by Suppressing Its MDM2-Mediated Ubiquitination. Biomol Ther (Seoul) 25 (4):396–403. doi:10.4062/biomolther.2017.086. http://www.ncbi.nlm.nih.gov/pubmed/28605833

Kanehisa, M., M. Araki, S. Goto, M. Hattori, M. Hirakawa et al., 2008KEGG for linking genomes to life and the environment. Nucleic Acids Res 36 (Database issue):D480–484. doi:10.1093/nar/gkm882. http://www.ncbi.nlm.nih.gov/pubmed/18077471

Kim, D., G. Pertea, C. Trapnell, H. Pimentel, R. Kelley et al., 2013 TopHat2: accurate alignment of transcriptomes in the presence of insertions, deletions and gene fusions. Genome Biol 14 (4):R36. doi:10.1186/gb-2013-14-4-r36. http://www.ncbi.nlm.nih.gov/pubmed/23618408

Langmead, B., C. Trapnell, M. Pop, and S.L. Salzberg, 2009 Ultrafast and memory-efficient alignment of short DNA sequences to the human genome. Genome Biol 10 (3):R25. doi:10.1186/gb-2009-10-3-r25. http://www.ncbi.nlm.nih.gov/pubmed/19261174

Lee, J.H., Y. Kang, V. Khare, Z.Y. Jin, M.Y. Kang et al., 2010 The p53-inducible gene 3 (PIG3) contributes to early cellular response to DNA damage. Oncogene 29 (10):1431–1450. doi:10.1038/onc.2009.438. http://www.ncbi.nlm.nih.gov/pubmed/20023697

Lee, S.U., and T. Maeda, 2012 POK/ZBTB proteins: an emerging family of proteins that regulate lymphoid development and function. Immunol Rev 247 (1):107–119. doi:10.1111/j.1600-065X.2012.01116.x. http://www.ncbi.nlm.nih.gov/pubmed/22500835

Leger, B., R. Cartoni, M. Praz, S. Lamon, O. Deriaz et al., 2006 Akt signalling through GSK-3beta, mTOR and Foxo1 is involved in human skeletal muscle hypertrophy and atrophy. J Physiol 576 (Pt 3):923–933. doi:10.1113/jphysiol.2006.116715. http://www.ncbi.nlm.nih.gov/pubmed/16916907

Li, B., N. Fillmore, Y. Bai, M. Collins, J.A. Thomson et al., 2014 Evaluation of de novo transcriptome assemblies from RNA-Seq data. Genome Biol 15 (12):553. doi:10.1186/s13059-014-0553-5. http://www.ncbi.nlm.nih.gov/pubmed/25608678

Liu, Q., C. Xu, S. Kirubakaran, X. Zhang, W. Hur et al., 2013 Characterization of Torin2, an ATP-competitive inhibitor of mTOR, ATM, and ATR. Cancer Res 73 (8):2574–2586. doi:10.1158/0008-5472.CAN-12-1702. http://www.ncbi.nlm.nih.gov/pubmed/234368011

Mao, X., T. Cai, J.G. Olyarchuk, and L. Wei, 2005 Automated genome annotation and pathway identification using the KEGG Orthology (KO) as a controlled vocabulary. Bioinformatics 21 (19):3787–3793. doi:10.1093/bioinformatics/bti430. http://www.ncbi.nlm.nih.gov/pubmed/15817693

Matsuda, E., O. Yu, and M. Kawaichi, 2008 CIBZ,a BTB-containing Zinc Finger Protein,Plays a Role in Apoptosis and Tumorigenesis in bit life sciences’ 1st annual world cancer congess-2008.

Matsuoka, S., B.A. Ballif, A. Smogorzewska, E.R. McDonald, 3rd, K.E. Hurov et al., 2007 ATM and ATR substrate analysis reveals extensive protein networks responsive to DNA damage. Science 316 (5828): 1160–1166. doi:10.1126/science.1140321. http://www.ncbi.nlm.nih.gov/pubmed/17525332

McGettigan, P.A., 2013 Transcriptomics in the RNA-seq era. Curr Opin Chem Biol 17 (1):4–11. doi:10.1016/j.cbpa.2012.12.008. http://www.ncbi.nlm.nih.gov/pubmed/23290152

Nishii, T., Y. Oikawa, Y. Ishida, M. Kawaichi, and E. Matsuda, 2012 CtBP-interacting BTB zinc finger protein (CIBZ) promotes proliferation and G1/S transition in embryonic stem cells via Nanog. Journal of Biological Chemistry 287 (15):12417. doi:10.1074/jbc.M111.333856. https://www.ncbi.nlm.nih.gov/pubmed/22315219

Reimann, E., S. Koks, X.D. Ho, K. Maasalu, and A. Martson, 2014 Whole exome sequencing of a single osteosarcoma case--integrative analysis with whole transcriptome RNA-seq data. Hum Genomics 8:20. doi:10.1186/s40246-014-0020-0. http://www.ncbi.nlm.nih.gov/pubmed/25496518

Robinson, M.D., D.J. McCarthy, and G.K. Smyth, 2010 edgeR: a Bioconductor package for differential expression analysis of digital gene expression data. Bioinformatics 26 (1):139–140. doi:10.1093/bioinformatics/btp616. http://www.ncbi.nlm.nih.gov/pubmed/19910308

Sasai, N., E. Matsuda, E. Sarashina, Y. Ishida, and M. Kawaichi, 2005 Identification of a novel BTB-zinc finger transcriptional repressor, CIBZ, that interacts with CtBP corepressor. Genes to Cells 10 (9):871–885. doi:10.1111/j.1365-2443.2005.00885.x. https://www.ncbi.nlm.nih.gov/pubmed/16115196

Sawai, C.M., J. Freund, P. Oh, D. Ndiaye-Lobry, J.C. Bretz et al., 2012 Therapeutic targeting of the cyclin D3:CDK4/6 complex in T cell leukemia. Cancer Cell 22 (4):452–465. doi:10.1016/j.ccr.2012.09.016. http://www.ncbi.nlm.nih.gov/pubmed/23079656

Shangary, S., and S. Wang, 2009 Small-molecule inhibitors of the MDM2-p53 protein-protein interaction to reactivate p53 function: a novel approach for cancer therapy. Annu Rev Pharmacol Toxicol 49:223–241. doi:10.1146/annurev.pharmtox.48.113006.094723. http://www.ncbi.nlm.nih.gov/pubmed/18834305

Shi, Y., X. Gao, Q. Hu, X. Li, J. Xu et al., 2016 PIK3C2A is a gene-specific target of microRNA-518a-5p in imatinib mesylate-resistant gastrointestinal stromal tumor. Lab Invest 96 (6):652–660. doi:10.1038/labinvest.2015.157. http://www.ncbi.nlm.nih.gov/pubmed/26950487

Stogios, P.J., G.S. Downs, J.J. Jauhal, S.K. Nandra, and G.G. Prive, 2005 Sequence and structural analysis of BTB domain proteins. Genome Biol 6 (10):R82. doi:10.1186/gb-2005-6-10-r82. http://www.ncbi.nlm.nih.gov/pubmed/16207353

The, I., S. Ruijtenberg, B.P. Bouchet, A. Cristobal, M.B. Prinsen et al., 2015 Rb and FZR1/Cdh1 determine CDK4/6-cyclin D requirement in C. elegans and human cancer cells. Nat Commun 6:5906. doi:10.1038/ncomms6906. http://www.ncbi.nlm.nih.gov/pubmed/25562820

VanArsdale, T., C. Boshoff, K.T. Arndt, and R.T. Abraham, 2015 Molecular Pathways: Targeting the Cyclin D-CDK4/6 Axis for Cancer Treatment. Clin Cancer Res 21 (13):2905–2910. doi:10.1158/1078-0432.CCR-14-0816. http://www.ncbi.nlm.nih.gov/pubmed/25941111http://www.ncbi.nlm.nih.gov/pubmed/25941111

Wang, D., M.A. Olman, J. Stewart, Jr., R. Tipps, P. Huang et al., 2011 Downregulation of FIP200 induces apoptosis of glioblastoma cells and microvascular endothelial cells by enhancing Pyk2 activity. PLoS One 6 (5):e19629. doi:10.1371/journal.pone.0019629. http://www.ncbi.nlm.nih.gov/pubmed/21602932

Wang, Z., M. Gerstein, and M. Snyder, 2009 RNA-Seq: a revolutionary tool for transcriptomics. Nat Rev Genet 10 (1):57–63. doi:10.1038/nrg2484. http://www.ncbi.nlm.nih.gov/pubmed/19015660

Wei, H., B. Gan, X. Wu, and J.L. Guan, 2009 Inactivation of FIP200 leads to inflammatory skin disorder, but not tumorigenesis, in conditional knock-out mouse models. J Biol Chem 284 (9):6004–6013. doi:10.1074/jbc.M806375200. http://www.ncbi.nlm.nih.gov/pubmed/19106106

Young, M.D., M.J. Wakefield, G.K. Smyth, and A. Oshlack, 2010 Gene ontology analysis for RNA-seq: accounting for selection bias. Genome Biol 11 (2):R14. doi:10.1186/gb-2010-11-2-r14. http://www.ncbi.nlm.nih.gov/pubmed/20132535

Zhang, Z., J. Chen, F. Chen, D. Yu, R. Li et al., 2017 Tauroursodeoxycholic acid alleviates secondary injury in the spinal cord via up-regulation of CIBZ gene. Cell Stress Chaperones. doi:10.1007/s12192-017-0862-1. http://www.ncbi.nlm.nih.gov/pubmed/29151236

Zhao, Q.Y., Y. Wang, Y.M. Kong, D. Luo, X. Li et al., 2011 Optimizing de novo transcriptome assembly from short-read RNA-Seq data: a comparative study. BMC Bioinformatics 12 Suppl 14:S2. doi:10.1186/1471-2105-12-S14-S2. http://www.ncbi.nlm.nih.gov/pubmed/22373417

